# Decodanda: a Python toolbox for best-practice decoding and geometric analysis of neural representations

**DOI:** 10.64898/2026.03.16.711920

**Authors:** Lorenzo Posani

**Author notes:** Correspondence: **Lorenzo Posani**.

## Abstract

Neural decoding is a powerful approach for inferring which variables are represented in the activity of a population of neurons, with broad applications ranging from basic neuroscience to clinical settings such as brain-computer interfaces. More recently, decoding has also been used as a cross-validated tool for studying the computationally relevant properties of representational geometry, revealing not only whether a variable is encoded, but also how it is encoded and which computations the collective activity of neural populations may support. However, decoding analyses present several technical challenges and common pitfalls that can lead to misleading conclusions if not handled carefully. Here, we introduce Decodanda, a Python toolbox for decoding and geometric analysis of neural population activity. Decodanda provides functions for decoding arbitrary variables and for quantifying geometric features of neural representations, including shattering dimensionality and cross-condition generalization performance (CCGP). Importantly, the package automates several essential best-practice safeguards, including trial-based cross-validation to avoid training-testing leakage from temporally correlated neural traces (a particularly important issue for calcium imaging data), null models for statistical significance, pseudo-population pooling, and cross-variable balancing to determine which of a set of correlated variables is genuinely encoded in the activity. Decodanda is agnostic to the specific classifier used for decoding, and it is designed to be both user-friendly and highly customizable, allowing researchers to assemble flexible analysis pipelines from modular building blocks. Here, we provide an overview of the design principles of Decodanda and illustrate its use cases in neuroscience research. Documentation, example notebooks, and source code are available at github.com/lposani/decodanda.

## Introduction

Recent breakthroughs in large-scale neural recording technologies have created an increasing demand for computational methods to investigate the structure of the neural code (1, 2). The term *neural decoding* encompasses a wide range of approaches, from linear regression to complex machine-learning methods (3), with the common goal of determining whether and how neural activity encodes information about external variables or the subject’s internal state. Because of its broad applicability, from basic research in animal models to clinical applications, decoding has become a standard tool in neuroscience research (3, 4).

In basic research, a typical application of neural decoding is to quantify the extent to which a neural population represents experimental or decision-related variables, such as spatial variables in the hippocampus(5–7), movement or sensory information in cortical areas (8–10), or higher-order cognitive variables such as context and rules in the prefrontal areas (11–14). Importantly, decoding can provide insight not only into whether variables are encoded in the collective activity of a neural population, but also into the structure in which they are encoded (15). A recent line of work has exploited this idea by using linear classifiers to probe the parallelism of coding directions across variables, linking geometric properties of neural representations to cognitive functions such as abstraction and generalization (13, 16), memory (17), decision-making (18), and inferential reasoning (14). Finally, decoding has also played a central role in clinical applications, including neural prostheses and brain-guided speech synthesis (19–21).

Recent methodological work on neural decoding has focused on improving predictive performance (3), developing unsupervised approaches to relate neural activity to external variables (22, 23), and lowering the technical barrier to entry through user-friendly software interfaces (24, 25). However, many of the most consequential pitfalls in neural decoding do not arise from the choice of classifier itself, but from how the data are organized, split, and balanced for the analysis (26).

If not properly addressed, these issues can lead to both false positives and false negatives. Common examples include inappropriate data segmentation for cross-validation, training-testing leakage due to temporal autocorrelation in slowly varying signals (such as calcium imaging or fMRI), null models that disrupt the data structure too strongly, making them too easy to outperform, and correlations among experimental variables that can act as confounds.

Decodanda is a Python package designed to implement best-practice safeguards against these pitfalls while exposing simple and customizable functions for decoding neural activity and analyzing representational geometry. Here, we describe its design principles and show how it can help researchers interrogate high-dimensional neural data sets.

### Installation

Decodanda can be installed via pip using

~~~
pip install decodanda
~~~

or by downloading the code from the repository: github.com/lposani/decodanda.

### Implementation: basics

Decodanda organizes neural data, or more generally feature vectors, into conditioned activity matrices corresponding to all combinations of values of the variables specified by the user. In the analysis pipelines (e.g., decoding, CCGP), Decodanda samples from these conditioned activity matrices to construct training and test sets according to the specific analysis to be performed. An example of this process for two binary variables is schematized in Fig. 1. The main inputs to the Decodanda constructor are two Python dictionaries: one containing the data (activations and labels), and the other containing the variables to be decoded.

**Fig. 1.**
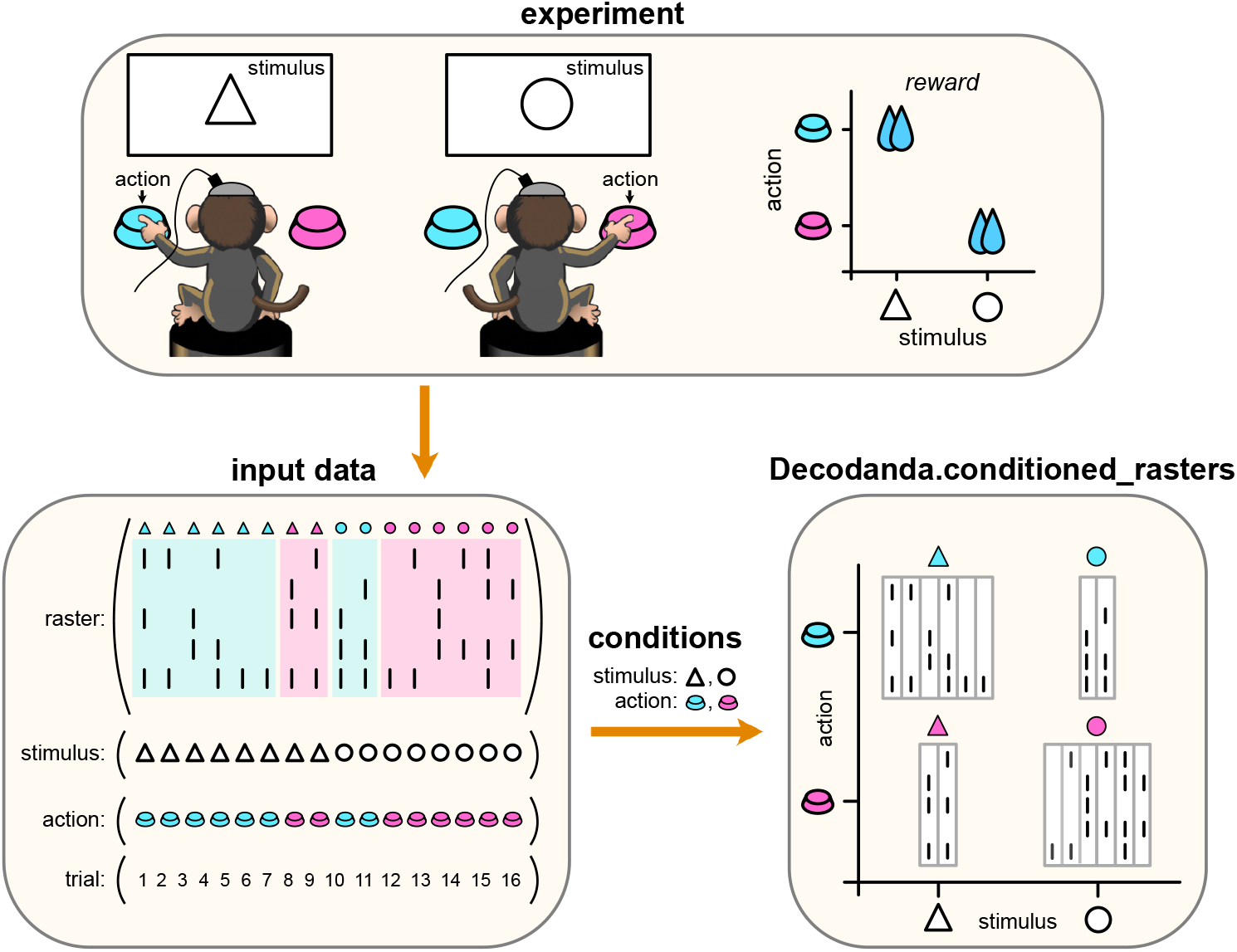
Data structure and basic design of Decodanda. This schematic shows how the Decodanda constructor manages data from a hypothetical experiment where 5 neurons are simultaneously recorded while a subject learns a stimulus-action association (triangle->blue button; circle->pink button, top panel). By passing a data structure with synchronized activity, variables, and trial number (bottom left panel) and a `conditions` dictionary that specifies what experimental variables we want to analyze, `Decodanda` will divide the neural activity matrix based on the four possible combinations of the two experimental variables (`stimulus`: triangle, circle; `action`: pink button, blue button) and store them in the `conditioned_rasters` class member (bottom right panel). These four conditioned activity matrices will then be used in the decoding and geometric analyses to investigate how the neural activity responds to the experimental conditions.

#### Input structure: data dictionary

For `N` recorded neurons and `T` trials (or time bins), the data dictionary must contain at least three entries corresponding to (1) a `TxN` neural activity matrix, (2) a `Tx1` labels/values vector for each variable we want to decode (e.g., behavior or stimuli), and (3) a `Tx1` trial index vector that will be used for cross-validation. A properly-formatted data set with two experimental variables (here, `stimulus and action`) would look like this (see also the Input Data panel in Fig. 1)

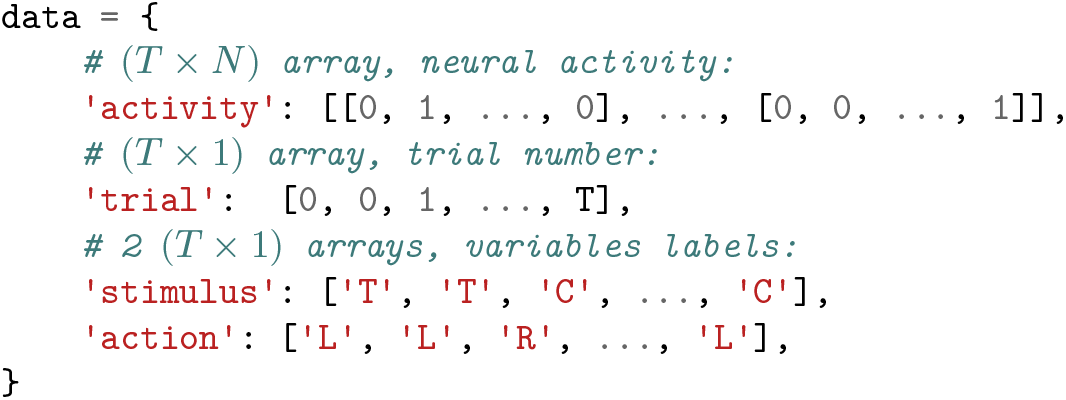

- `data[‘activity’]` is the set of features we want to analyze. Each feature can be continuous (e.g., calcium fluorescence) or discrete (e.g., spikes) values. The features have to be organized into a `TxN` array under the `activity`key.
- `data[‘stimulus’]`and `data[‘action’]` are the values of the variables we want to decode from the neural features. Each variable has to be structured into a `Tx1` array synchronized with the neural activity matrix.
- `data[‘trial’]` is a `Tx1` array that specifies which chunks of the data are to be considered statistically independent for the purpose of cross-validations (more on this below).

#### Input structure: conditions dictionary

The input data structure can contain any number of variables with any number of values, but only those specified in the `conditions` dictionary are used in the analysis:

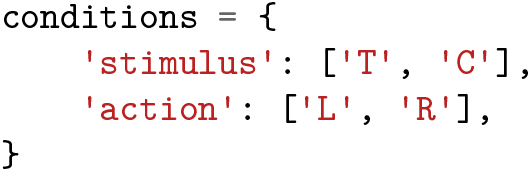

Decodanda defines conditions as the intersections of the specified variable values. These conditioned subsets form the basic units from which decoding and geometric analyses are constructed. This representation also makes it possible to balance across variables when needed, for example to disentangle correlated experimental factors. Users should therefore be parsimonious when defining the `conditions` dictionary. As the number of variables and allowed values increases, the total number of condition combinations grows combinatorially, and each relevant combination must be sufficiently represented in the data for reliable analysis.

#### Conditioned activity matrices

By constructing a `Decodanda`object from the `data` and `conditions` dictionaries,

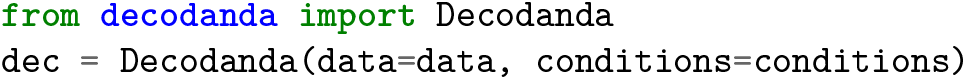

the neural activity is partitioned into conditioned activity matrices, one for each combination of the specified variable values. In the example above, this yields four conditioned subsets:`(stimulus = T, action = L), (stimulus = T, action = R), (stimulus = C, action = L), and (stimulus = C, action = R).`

#### Trial structure

The `trial` structure plays a central role in `Decodanda,` because it determines how samples are grouped into statistically independent units for cross-validation. Rather than splitting individual activity vectors independently, `Decodanda` uses the`trial` labels to ensure that all samples sharing the same trial identity are assigned together to either the training set or the test set. This prevents information from leaking across the train-test split when nearby samples are not statistically independent.

This is especially important when neural features are temporally correlated over timescales longer than the sampling interval, as is often the case for calcium imaging, fMRI, or slowly varying behavioral variables (26). In such cases, randomly splitting individual time points can produce artificially inflated decoding performance, because the decoder may exploit temporal proximity rather than similarity across independent observations.

In many experimental paradigms, the relevant trial structure is naturally defined by the task and can be passed directly through the `trial` entry of the data dictionary. In other situations, however, there may be no obvious trial segmentation, for example during free behavior recorded continuously in a single session. In these cases, the user should define a pseudo-trial structure that partitions the recording into chunks separated sufficiently in time to minimize temporal leakage across training and test sets.

As a convenience, `Decodanda` also supports a default pseudo-trial definition in which trials are identified as contiguous chunks of data that share the same values for all variables specified in the analysis. This behavior can be invoked by setting `trial=None` in the `Decodanda` constructor. However, this should be treated as a practical fallback, and users are encouraged to verify that it provides adequate temporal separation for their data and experimental design.

Finally, when an explicit time variable is available, `Decodanda` provides an additional safeguard against temporal leakage through the `min_time_separation` parameter. By specifying the time vector through `time_attr,` the user can require a minimum temporal separation between samples assigned to different trials. For example, assuming that the `data`dictionary contains a `time (s)`keyword, tracking time in seconds,

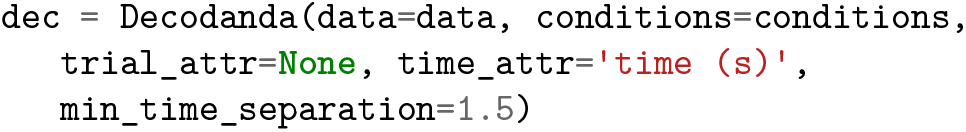

will create a pseudo-trial logic using contiguous time bins that share the same condition and discard the minimum amount of pseudo-trials to ensure that they are all at least separated by a second and a half. Note that this procedure can reduce the effective amount of data available for the analysis.

#### Choosing a classifier

The design of Decodanda makes it agnostic to the classifier used for decoding. By default, Decodanda uses a scikit (27) SVM with a linear kernel (sklearn.SVM.linearSVC). However, the user can specify any classifier in the Decodanda constructor, as long as it is clon-able (by sklearn.base.clone) and exposes the methods `fit, predict,` and `score.` Besides the vast choice of `sklearn` classifiers, the user can implement any personalized classifier by creating a child class of `sklearn.base.BaseEstimator.` See for example the [sci-kit documentation](https://scikit-learn.org/stable/developers/develop.html) on how to implement custom classifiers.

#### Specifying conditions using lambda functions

Sometimes conditions are more complex than a simple set of discrete labels. For example, one may wish to decode activity corresponding to selected intervals of a continuous variable, or to restrict the analysis using multiple criteria simultaneously. `Decodanda` therefore allows conditions to be specified by arbitrary Python callables, including lambda functions. This provides full flexibility in defining how samples are assigned to each condition, as long as the subsets corresponding to different values of the same variable are disjoint.

For example, we may want to decode action and stimulus from the data above while restricting the analysis to the first ten trials. Rather than adding a new field to the data dictionary, this can be expressed directly using lambda functions:

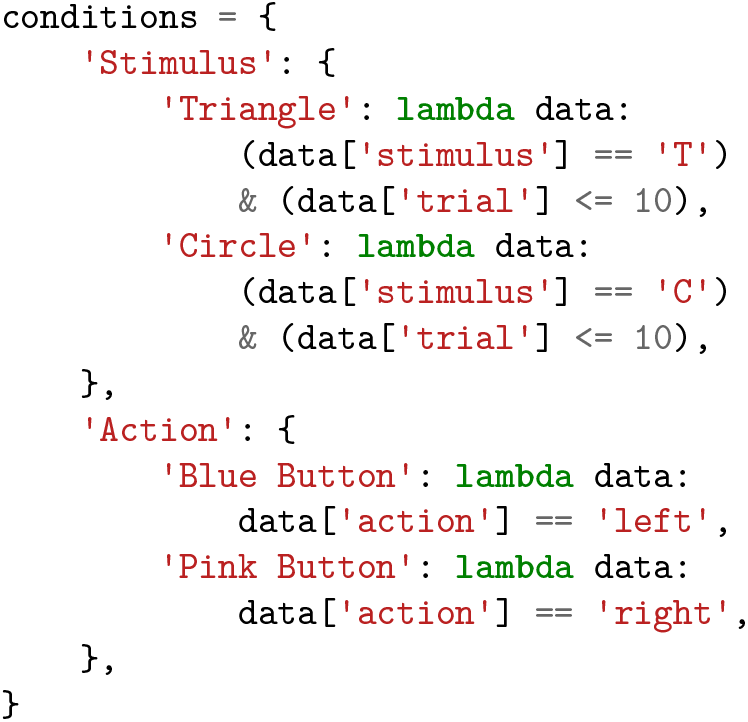

In this example, the restriction `data[‘trial’] <=`10 only needs to be specified once, in the definition of the `Stimulus` conditions. Since `Decodanda` constructs joint conditioned subsets by intersecting the conditions specified for all variables, the resulting subsets for `Action` are automatically restricted to the same trial range.

### Decoding

#### Cross-variable balancing

Experimental variables are often correlated with one another. Although well-controlled experiments are typically designed to minimize such confounds, more naturalistic settings, such as freely behaving subjects, or paradigms in which variables are correlated by design (for example, through stimulus-action associations), may not provide sufficient control to ensure balanced sampling across all relevant variables. This can lead to misleading decoding results, because successful decoding of one variable may in fact be driven by neural responses to another correlated variable. For example, in the task illustrated in Fig. 1, the action performed by a trained subject is correlated with the presented stimulus through reinforced learning. Without appropriate balancing, this correlation could allow `action` to be decoded from neural activity that in reality only reflects visual responses to the stimulus.

To address this problem, `Decodanda` implements cross-variable balanced sampling in its decoding procedures. Specifically, in all decoding pipelines, conditions (task-variable combinations) are resampled to ensure equal representation in the training and testing sets. This allows users to control for confounding variables by including them in the `conditions` dictionary, thereby testing whether a target variable can still be decoded once correlated factors have been balanced.

#### Cross-validation and trial structure

As specified above, a well-structured data set for `Decodanda` includes a `trial` index that specifies which subsets of the data can be treated as statistically independent samples of neural activity for a given condition. The `Decodanda` decoding methods (see [documentation](https://decodanda.readthedocs.io/en/latest/modules.html#decoding-functions)) use these trial labels to partition the data into training and testing sets. A schematic of the decoding pipeline for a variable is shown in Fig. 2. The procedure can be summarized in five main steps:

**Fig. 2.**
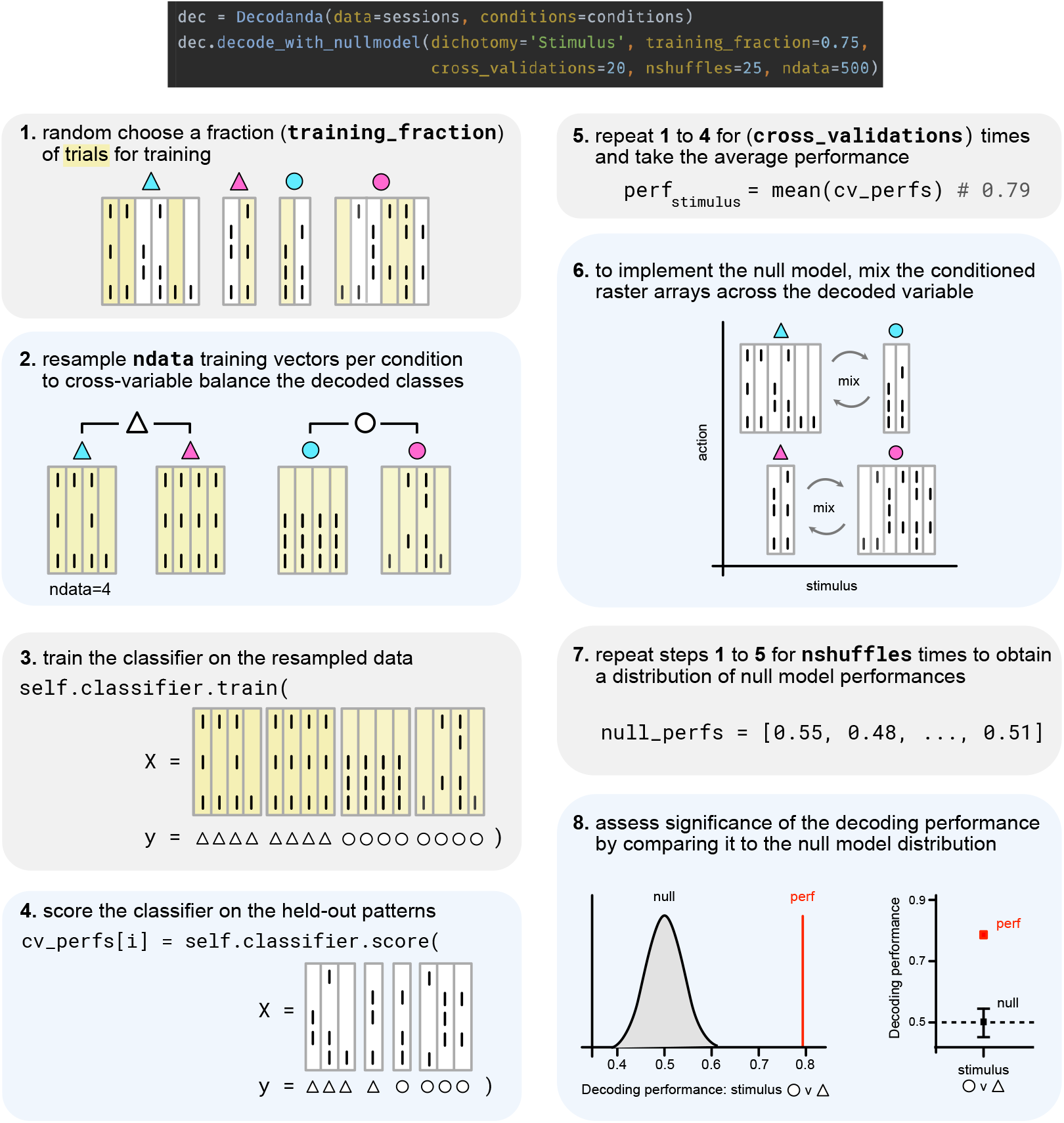
Schematic of the decoding pipeline implemented by Decodanda.decode: drawing from the example experimental setup in Fig. 1, the conditioned activity matrices for the four conditions (triangle-blue, triangle-pink, circle-blue, circle-pink) are shown in panel (1). In this example, individual population vectors of 5 neurons are sampled on independent trials. Training trials are highlighted in yellow. The figure shows the individual steps performed by the `Decodanda.decode` function for the variable `stimulus` (triangle vs. circle). See the main text for a detailed explanation.

1. Each conditioned activity matrix is partitioned into training and testing trials using the `trial` index, ensuring that activity vectors sharing the same trial label are always assigned together to either the training or the testing set. The fraction of trials assigned to training is controlled by the `training_fraction` argument.
2. The same number of activity vectors is then sampled, with replacement, from the training portion of each conditioned activity matrix and concatenated to form the training array. This balanced re-sampling ensures that non-decoded variables remain matched across conditions, thereby reducing confounding effects due to unbalanced or correlated variables. The number of sampled activity vectors is controlled by the `ndata` argument.
3. The classifier is trained on the sampled training data. Any classifier can be used, provided that it exposes the expected interface (see the section “Choosing a classifier”).
4. The classifier is then evaluated on the held-out test data, i.e., activity vectors that were not used for training, yielding a decoding performance for that train-test split.
5. Steps 1 to 4 are repeated multiple times, as controlled by the `cross_validations` argument, to obtain a distribution of decoding performances across different random training-test partitions. The mean across these iterations is reported as the decoding performance on the data.

#### Null model

A single decoding performance value - such as the one returned by the `Decodanda.decode` function - can be difficult to interpret without an appropriate reference. To place this value in context, `Decodanda` implements a null-model procedure that generates a distribution of decoding performances compatible with the null hypothesis that labels (e.g., behavior) and features (e.g., spikes) are independent. This allows the significance of the observed decoding performance to be assessed by assigning it a *p*-value, i.e., the probability of observing an equal or greater performance under the null hypothesis. Importantly, the null model is implemented *upstream* of the decoding pipeline described above, so that the structure of the original data is preserved as much as possible except for the relationship between labels and features. The null-model procedure can be summarized in three main steps (numbered 6 to 8 in Fig. 2):

1. The first step is to break the relationship between labels and features by randomly shuffling the assignment of activity vectors across conditions. If, for example, the decoded variable is `stimulus,` the procedure will mix activity from conditions such as`(stimulus=T, action=L) and (stimulus=C, action=L).` This shuffling is performed subject to three constraints:
  - When possible, condition assignments are shuffled only across the decoded variable, while preserving the structure of the other variables.
  - The shuffling is compatible with the trial structure used for cross-validation, so that all activity vectors sharing the same trial label remain grouped together throughout the procedure.
  - The total number of samples assigned to each condition is preserved.
2. The cross-validated decoding procedure described in the previous section and in Fig. 2 (steps 1 to 5) is then applied to this shuffled version of the data, yielding one null-model decoding performance value.
3. The shuffling and decoding steps are repeated the number of times specified by the keyword `nshuffles,` producing a null distribution of decoding performance values. The significance of the observed decoding performance can then be quantified either using a Gaussian approximation based on the null distribution `(ptype=‘z’,` default behavior) or by computing the empirical fraction of null-model values that exceed the observed value `(ptype=‘count’).`

### Cross-Condition Generalization Performance

Cross-condition generalization performance (CCGP) is a geometric metric that quantifies how well a variable can be decoded across changes in the other variables that define the experimental conditions (13). High CCGP indicates that the coding direction associated with the decoded variable is similar across different conditions, i.e., that it generalizes beyond the specific combinations of the remaining variables used for training. In the ideal case, a set of variables with high CCGP is associated with an approximately parallel and low-dimensional geometric organization of the neural representations, akin to a cuboid structure in neural activity space (13). More generally, CCGP is an example of how geometric properties of neural representations - in this case, the parallelism of coding directions -can be related to cognitive functions such as generalization and abstraction. This relation has been explored in recent studies across different species and tasks (13, 14, 16–18, 28, 29).

`Decodanda` implements CCGP analysis as a particular cross-condition decoding procedure using a linear classifier: the classifier is trained to decode a chosen variable from a subset of conditions and then tested on held-out conditions that differ in the values of the non-decoded variables. The resulting performance quantifies how well the coding strategy learned in one subset of conditions generalizes to new ones. Figure 3 outlines the detailed steps followed by `Decodanda` to compute CCGP (see also the [documentation](https://decodanda.readthedocs.io/en/latest/modules.html#geometry-functions)):

**Fig. 3.**
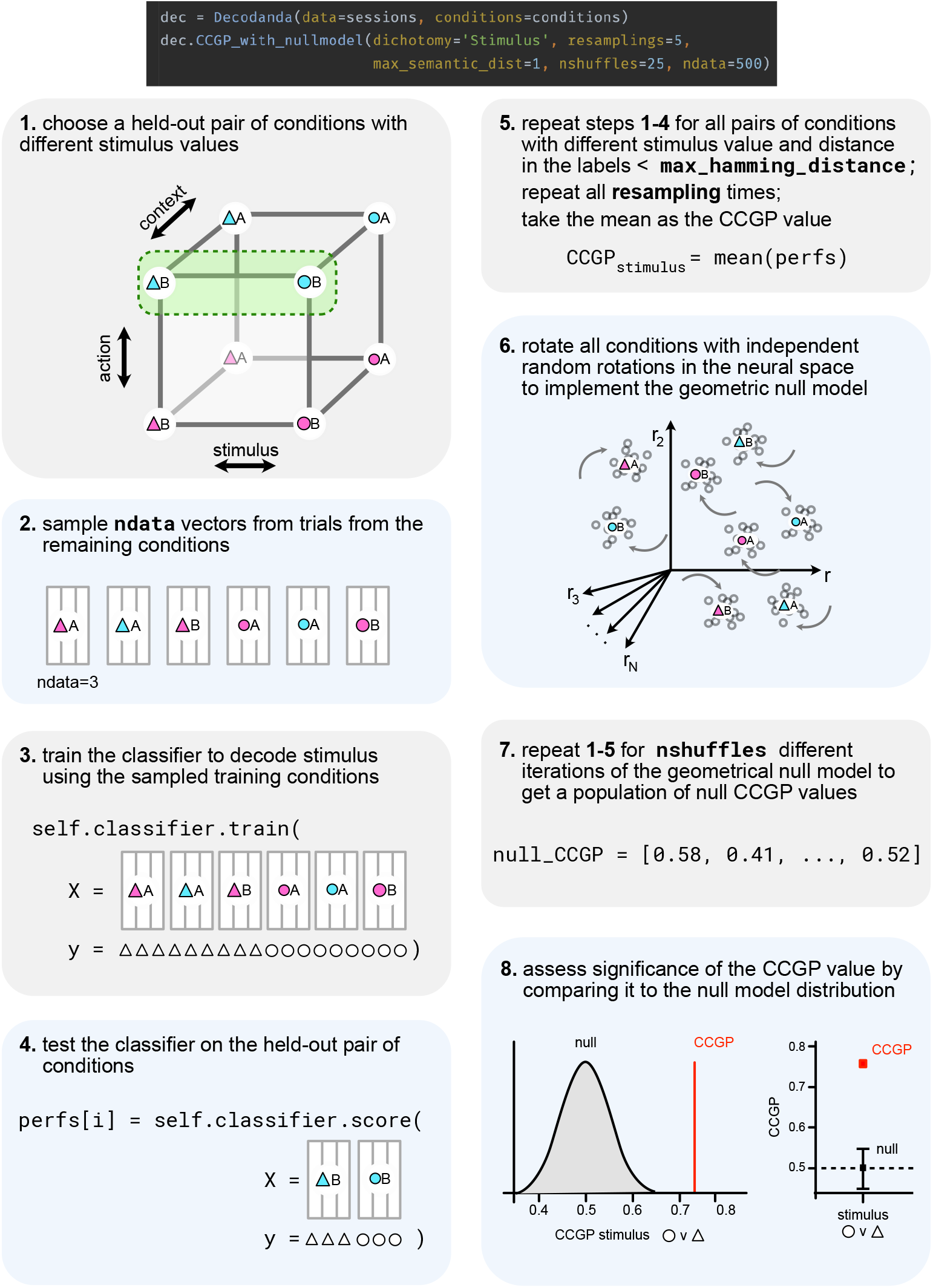
Schematic of CCGP protocol: The classifier is trained on a set of conditions and then tested on different, held-out conditions to assess generalization. The CCGP score is the average performance across all condition pairs.

1. A pair of conditions belonging to opposite sides of the decoded dichotomy is selected to define the held-out test set. The admissible held-out pairs are determined by the `max_semantic_dist` argument, which specifies the maximum semantic distance between the two test conditions, i.e., the maximum number of variables whose values are allowed to differ between them. For example, with two variables and `max_semantic_dist=1,` the pair `(T, L)–(C, L)` can be used as a held-out test pair when decoding `stimulus,` whereas `(T, L)–(C, R)` is excluded because both `stimulus` and `action` change.
2. A given number `(ndata)` of activity vectors is then sampled from each of the remaining conditions. As in standard decoding, this balanced sampling ensures that the classifier is not biased by unequal representation of the other variables.
3. The sampled activity vectors are used to train the classifier to decode the chosen variable (e.g., `stimulus`).
4. The classifier is then evaluated on the held-out pair of conditions, measuring how well decoding of the chosen variable generalizes to a new combination of the non-decoded variables.
5. Steps 1 to 4 are repeated over all admissible held-out condition pairs, as determined by `max_semantic_dist.`

The final CCGP score is the mean decoding performance across these cross-condition train-test splits.

#### Geometric null model

As with standard decoding, the absolute value of CCGP may be difficult to interpret without an appropriate reference. `Decodanda` therefore implements a geometric null model that disrupts the relationships between conditions while preserving their decodability (13). This allows the user to compare the observed CCGP score with a null distribution and to assess whether the observed degree of cross-condition generalization exceeds what would be expected in the absence of a geometric structure. As for standard decoding, the significance of the observed CCGP can be quantified either using a Gaussian approximation based on the null distribution or by computing the empirical fraction of null values that exceed the observed score.

1. For each null-model iteration, all conditions are transformed by random, independent rotations in neural activity space. This disrupts the geometric relationships between conditions while preserving the identity of each condition and its within-condition covariance structure.
2. The CCGP procedure described above is then repeated on this transformed data for the number of iterations specified by `nshuffles`, producing a null distribution of CCGP values.
3. Finally, the observed CCGP score is compared against this null distribution, analogously to the decoding analysis, to assess its statistical significance.

### Pseudo-population pooling from multiple data sets

`Decodanda` facilitates the pooling of data from multiple recording sessions or subjects into a single pseudo-population. This feature is useful for combining neurons recorded in different experiments or individuals into a common decoding analysis, thereby increasing the dimensionality of the population and exploiting the combinatorial sampling of activity vectors across data sets. The resulting pseudo-population vectors have dimensionality equal to the total number of recorded neurons across sessions or subjects.

Pseudo-population pooling is straightforward to invoke in `Decodanda,` by simply passing a list of datasets to the`data` argument:

~~~
dec = Decodanda(data=[data1, data2, …],
conditions=conditions,
**decodanda_params)
~~~

Operationally, pseudo-populations are constructed by first resampling condition-matched activity vectors within each individual data set and then concatenating those sampled vectors across data sets (Fig. 4). The resulting pseudo-population can then be used for decoding and geometric analyses.

**Fig. 4.**
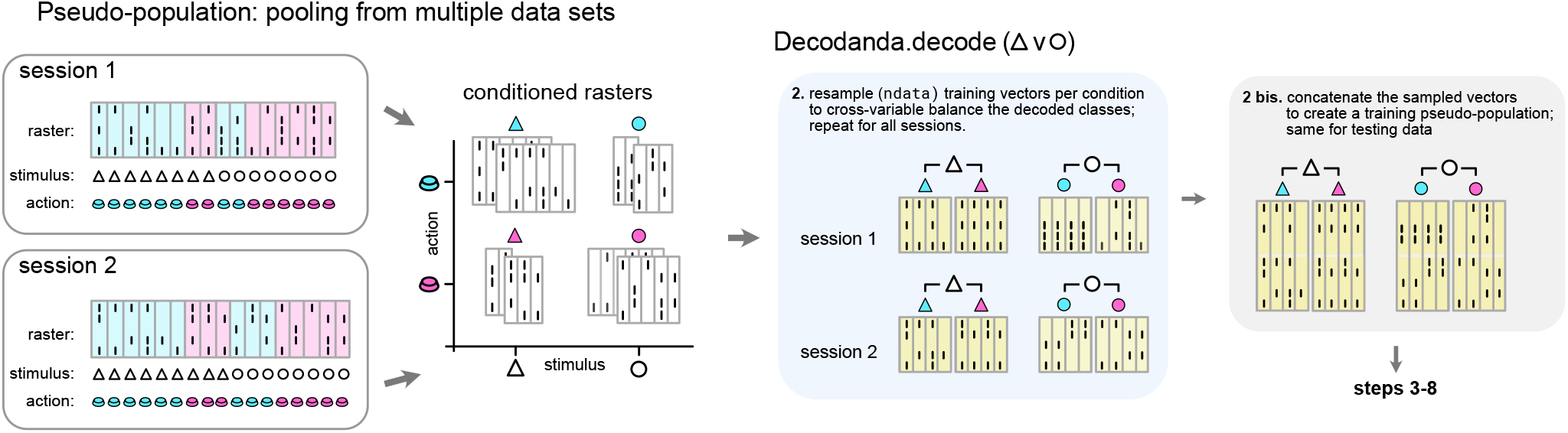
Pseudo-population pooling across multiple data sets. **(a)** Example of two independent sessions with different numbers of recorded neurons. **(b)** Each session is divided into conditioned activity matrices. **(c)** Training and testing splits are performed separately in each session, and balanced activity vectors are resampled within each condition. **(d)** Resampled activity vectors from different sessions are concatenated to form pseudo-population vectors, which are then used for decoding.

#### Caveats

When decoding from pseudo-population vectors, a few important points should be kept in mind:

- **Cross-validation**: Every training-testing split must be performed *before* resampling and concatenating activity vectors into a pseudo-population. This is essential to ensure that the same data are not resampled into both training and testing sets. `Decodanda` implements this procedure automatically.
- **CCGP and geometry**: Geometric results obtained from pseudo-populations should be interpreted with caution, since concatenating activity across sessions or subjects can substantially change the representational geometry. A single subject or session may drive the decoding or geometric structure of the pooled population. Pseudo-population results should therefore always be compared with the corresponding analyses in individual subjects whenever possible.
- **Noise correlations**: Pseudo-populations are often constructed by sampling single neurons independently across sessions. This approach has been criticized because it destroys the covariance structure of simultaneously recorded activity, including noise correlations, which can alter decoding performance (7, 30). `Decodanda` avoids this issue by resampling whole population vectors of simultaneously recorded neurons, rather than sampling individual neurons independently.

### Interpreting the decoding and CCGP results

Different representational geometries have important implications for the computational properties of the neural code (13, 17). Using the `decode` and `CCGP` functions in `Decodanda,`one can adopt an inverse approach and infer aspects of representational geometry directly from the computational properties of neural recordings.

The key idea is that decoding and CCGP provide complementary information. Standard decoding quantifies whether a variable is linearly accessible from population activity, i.e., whether it can be read out by a classifier. CCGP instead quantifies whether this readout generalizes across changes in the other variables that define the task conditions. A variable may therefore be decodable without being abstract: in that case, the neural code contains information about the variable, but in a condition-specific format that does not generalize well across contexts. Conversely, high decoding together with high CCGP indicates that the variable is represented in a format that is both separable and approximately invariant across the remaining task dimensions.

In Fig. 5, we show three exemplar geometries for two variables, `stimulus` and `action,` together with their corresponding decoding fingerprints, i.e., the combination of decoding and CCGP values expected from each geometry. These examples illustrate that similar decoding values can arise from geometrically distinct representations, and that combining decoding with CCGP allows one to distinguish between them. For a more complete discussion of the geometric interpretation of these metrics, together with practical coding examples, see the following [notebook material](http://www.tinyurl.com/attn-geometry) from the 2024 graduate course in Advanced Topics in Theoretical Neuroscience at Columbia University.

**Fig. 5.**
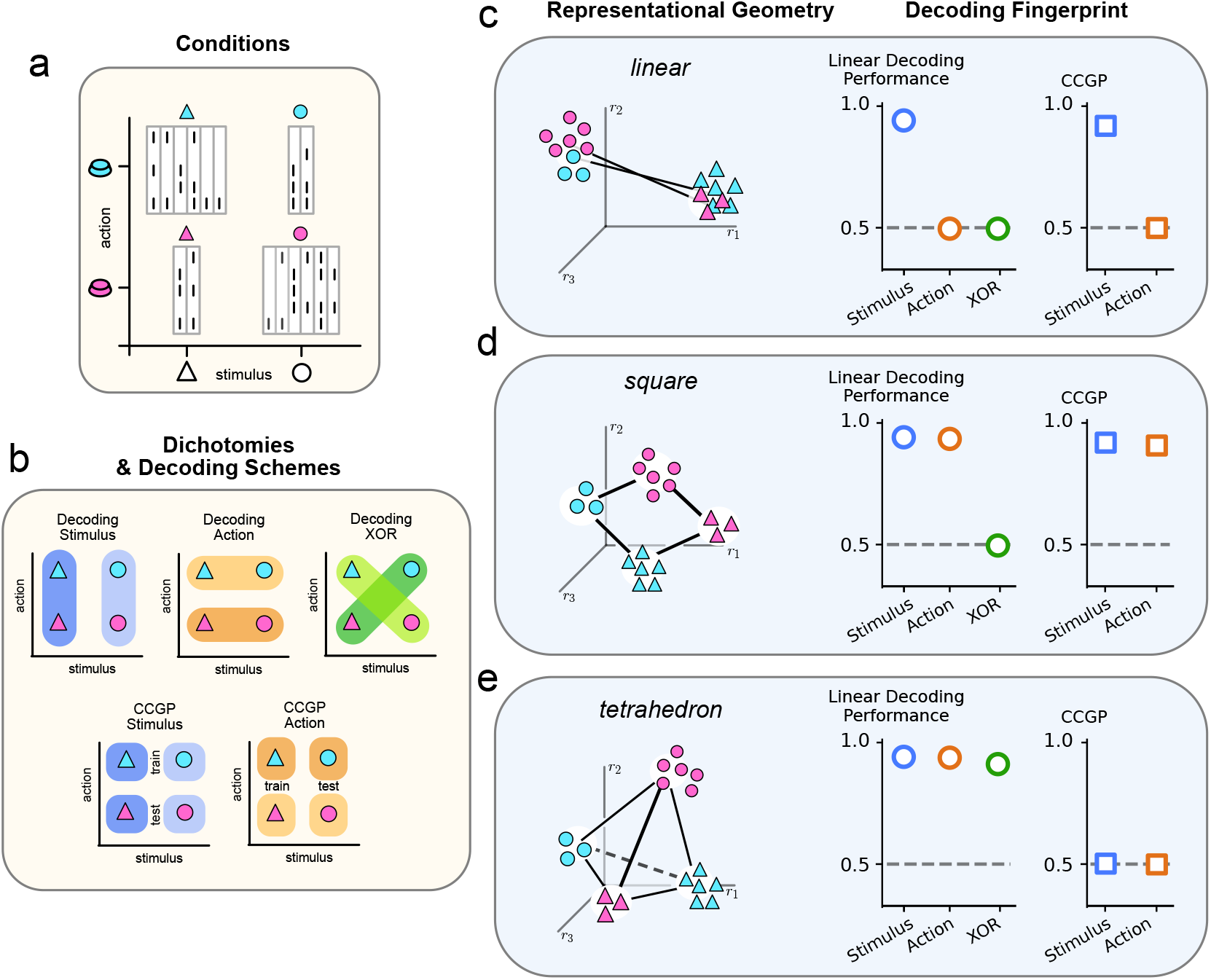
Interpretation of decoding results. **(a)** unbalanced conditions separated into `conditioned_rasters` from the Decodanda class (from the example of Fig. 1). **(b)** Conceptual schemas of decoding and CCGP analyses for the two variables (Stimulus and Choice). **(c)** Schematic and decoding fingerprint (combination of decoding and CCGP results) of a linear geometry: one variable encoded and abstract. **(d)** Schematic and decoding fingerprint of a squared geometry: two variables encoded and abstract. **(c)** Schematic and decoding fingerprint of a high-dimensional geometry: two variables encoded, no abstract variables.

#### Shattering dimensionality

While decoding and CCGP focus on specific task variables, shattering dimensionality (SD) measures how many dichotomies of the experimental conditions can be linearly separated by the neural activity (12, 13). High shattering dimensionality indicates that the population representation is sufficiently rich to support many distinct linear readouts, and has been linked to representational flexibility and memory capacity (12, 17). On the contrary, lower shattering dimensionality is consistent with a more constrained and low-dimensional organization.

Decodanda computes SD by performing a cross-validated decoding analysis over all possible balanced dichotomies of the set of conditions. Intuitively, this measures how many distinct balanced partitions of the condition space can be linearly separated by the neural activity. For a total of *N* conditions, the number of balanced dichotomies is given by 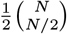 In the case of *n* binary variables, *N* = 2^*n*^, so this number grows extremely rapidly (for 4 binary variables, there are 6435 balanced dichotomies). Caution is therefore advised when analyzing more than three variables, or variables with more than two values, as the number of dichotomies can quickly become computationally prohibitive.

#### Documentation, Notebooks, and Real-World Applications

Additional information and practical examples of `Decodanda’s` main functionalities can be found in the [documentation](https://decodanda.readthedocs.io/en/latest/modules.html), the GitHub repository, and the accompanying [notebooks](https://github.com/lposani/decodanda/tree/master/notebooks). `Decodanda` has played a central methodological role in recent studies of neural representational geometry (17, 28, 31). It has also been used more broadly as a decoding tool in collaborative studies addressing a range of neuroscience questions (32–35).

## Acknowledgements

Decodanda is based on the collective knowledge and expertise of many colleagues in the Fusi Lab and in the Center for Theoretical Neuroscience. The author is especially grateful to Stefano Fusi for his continuous guidance and mentoring, and to M Rigotti, F Stefanini, M Benna, R Nogueira, V Fascianelli, J Minxha, S Muscinelli, and J Johnston for the numerous discussions that made this work possible. This work received support from NIH RF1AG080818, the Kavli Foundation, and the Gatsby Foundation GAT3708 and by the NIH 1K99MH135166-01 grant.

